# Emergence of behavior in a self-organized living matter network

**DOI:** 10.1101/2020.09.06.285080

**Authors:** Philipp Fleig, Mirna Kramar, Michael Wilczek, Karen Alim

## Abstract

What is the origin of behavior? Although typically associated with a nervous system, simple life forms also show complex behavior – thus serving as a model to study how behaviors emerge. Among them, the slime mold *Physarum polycephalum*, growing as a single giant cell, is renowned for its sophisticated behavior. Here, we show how locomotion and morphological adaptation behavior emerge from self-organized patterns of rhythmic contractions of the actomyosin lining of the tubes making up the network-shaped organism. We quantify the spatio-temporal contraction dynamics by decomposing experimentally recorded contraction patterns into spatial contraction modes. Surprisingly, we find a continuous spectrum of modes, as opposed to few dominant modes. Over time, activation of modes along this continuous spectrum is highly dynamic, resulting in contraction patterns of varying regularity. We show that regular patterns are associated with stereotyped behavior by triggering a behavioral response with a food stimulus. Furthermore, we demonstrate that the continuous spectrum of modes and the existence of irregular contraction patterns persist in specimens with a morphology as simple as a single tube. Our data suggests that the continuous spectrum of modes allows for dynamic transitions between a plethora of specific behaviors with transitions marked by highly irregular contraction states. By mapping specific behaviors to states of active contractions, we provide the basis to understand behavior’s complexity as a function of biomechanical dynamics. This perspective will likely stimulate bio-inspired design of soft robots with a similarly rich behavioral repertoire as *P. polycephalum*.

## Introduction

Survival in changing environments requires from organisms the ability to switch between diverse behaviors (1, 2). In higher organisms, rapid changes in neural activity enable this capacity, ranging from almost random to strongly correlated firing patterns of neurons (3, 4). However, many organisms without a nervous system are also able to readily transition between a multitude of behaviors, which suggests that the underlying biophysical processes display a dynamic variability analogous to that of a nervous system.

Examples of specific behaviors in non-neural organisms include the run-and-tumble chemotaxis of the *E. coli* bacterium (5), environmentally cued aggregation of the social amoeba *Dictyostelium discoideum* (6, 7) or the cooperative growth behavior of *B. subtilis* bacterial colonies (8, 9). An organism with an exceptionally versatile behavioral repertoire is the slime mould *Physarum polycephalum* – a unicellular, network-shaped organism (10) of macroscopic dimensions, typically from a millimeter to tens of centimeters.

*P. polycephalum’*s complex behavior is most impressively demonstrated by its ability to solve spatial optimization and decision-making problems (11–15), exhibit habituation to temporal stimuli (16), and use exploration versus exploitation strategy (17). The generation of such rich behavior requires a mechanism allowing not only for long-range spatial coordination but also the flexibility to enable switching between different specific behavioral states.

The behavior generating mechanism in *P. polycephalum* are the active, rhythmic, cross-sectional contractions of the actomyosin cortex lining the tube walls (18–20). The contractions drive cytoplasmic flows throughout the organism’s network (21, 22), transporting nutrients and signalling molecules (23). Cytoplasmic flow is responsible for mass transport across the organism and thereby contractions directly control locomotion behavior (24–28).

So far only one type of network-spanning peristaltic contraction pattern has been described experimentally (22, 29). However, for small *P. polycephalum* plasmodial fragments various other short-range contraction patterns have been observed (25, 26) and predicted by theory of active contractions (30–34). Similarly, up to now unknown complex, large-scale contraction patterns might play a role in generating the behavior of large *P. polycephalum* networks. Furthermore, transitions between such large-scale patterns are needed to allow for switching between specific behaviors.

Here, we decompose experimentally recorded contractions of a large *P. polycephalum* network of stable morphology into a set of physically interpretable contraction modes using Principal Component Analysis. Surprisingly, we find a continuous spectrum of modes and high variability in the activation of modes within this spectrum. By perturbing the network with an attractive stimulus, we show that the resulting locomotion response is coupled to a selective activation of regular contraction patterns. Guided by these observations, we design an experiment on a *P. polycephalum* specimen reduced in morphological complexity to a single tube. This allows us to quantify the causal relation between locomotion behavior, cytoplasmic flow rate and varying types of contraction patterns, thus revealing the central role of dynamical variability to generate different behaviors.

## Results

### Continuous spectrum of contraction modes reveals large variability in organism’s contraction dynamics

To characterize the contraction dynamics of a *P. polycephalum* network, we record contractions using bright-field microscopy and decompose this data into a set of modes using Principal Component Analysis (PCA). At first, networks in bright-field images are skeletonized, with every single skeleton pixel representing the local tube intensity as a measure of the local contraction state (35). Thus, any network state at a time *t_i_* is represented by a list of pixels, 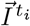, along the skeleton, see Fig. 1A. Performing PCA on this data results in a linear decomposition of the intensity vectors 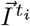 into a basis of modes 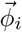 (SI Text):

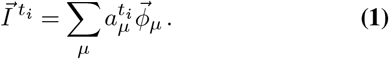

**Fig. 1.**
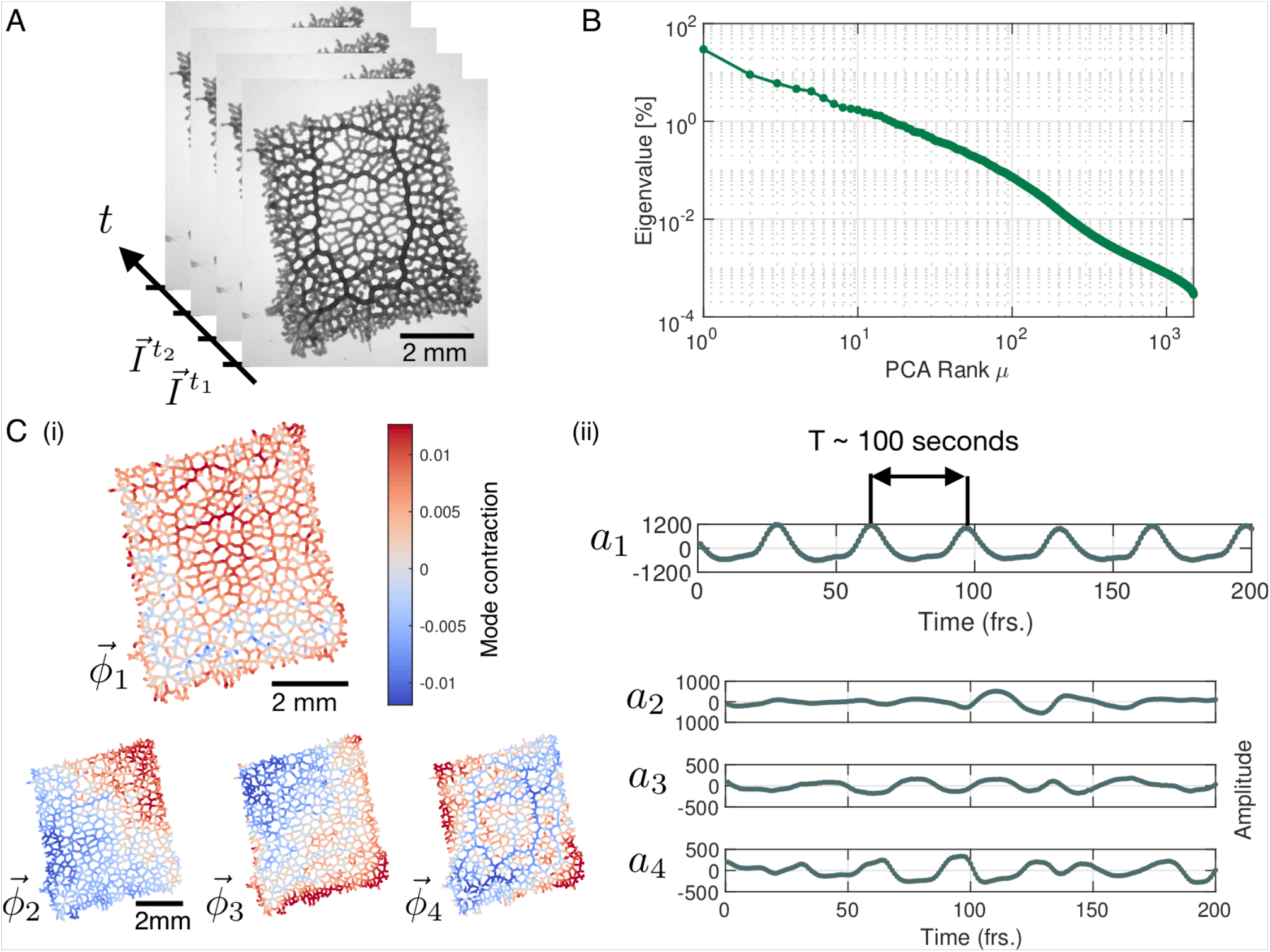
Principal Component Analysis yields a continuous spectrum of contraction modes in the *P. polycephalum* network. (A) Exemplary stack of bright-field images of the recorded network. Pixel intensities encode the contraction state (tube dilation) at each point of the network. Principal Component Analysis is performed on a stack of post-processed bright-field frames. (B) Relative eigenvalues in percent, plotted against the mode rank *μ* on a log-log graph. The eigenvalue spectrum is continuous, without a natural cutoff. (C) (i) Structure of the four highest-ranking modes 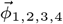 with their respective coefficients shown in (ii). The red-blue color spectrum indicates the contraction state. The modes are eigenvectors of the covariance matrix. The coefficient *a*_1_ of the first mode captures the organism’s characteristic oscillation period of ≈100sec, while the coefficients *a*_2,3,4_ show considerable variation in amplitude and frequency over time.

The modes, 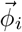 are orthonormal eigenvectors of the covariance matrix of the data and represent linearly uncorrelated contraction patterns of the network, and 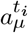 denotes the timedependent coefficients of the modes.

We rank modes according to the magnitude of their eigenvalue. Contrary to the small number of large eigenvalues found in a number of biological systems (36–38), here the spectrum of relative eigenvalues (SI Text) reveals a continuous distribution of eigenvalues with no natural cutoff Fig. 1B. As a result, PCA does not directly lead to a dimensionality reduction of the data. Instead, we here investigate the characteristics of mode dynamics that result from a continuous spectrum and how these shape the organism’s behavior.

The highest ranking modes shown in Fig. 1C(i) have a smooth spatial structure that varies on the scale of network size. As we will discuss below, such large-scale modes are associated with the long wavelength peristalsis observed in (21, 22). Interestingly, we also find modes highlighting specific morphological characteristics of the network. For example, the structure of mode 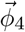, Fig. 1A, corresponds to the thickest tubes of the network, which suggests a special role of these tubes in the functioning of the network. Finally, as we go to lower ranked modes, the spatial structure of the modes becomes increasingly finer. Yet, despite lacking an obvious interpretation for their structures, like for mode 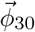, *SI Appendix* Fig. S1, it is not possible to ignore their contribution relative to high ranking modes.

Next, we turn to the time-dependent coefficients of modes shown in Fig. 1C(ii). In accordance with the known rhythmic contractions (39) the coefficient *a*_1_ of the highest ranked mode 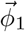 oscillates with a typical period of *T* ~ 100 sec. Most strikingly, amplitudes of mode coefficients vary significantly over time – even on orders of magnitude.

To map out the complexity of contractions over time, we define a set of *significant modes* for every time point. We quantify the activity of a mode by its *relative amplitude*

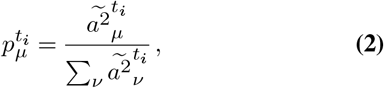

where 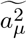 denotes the amplitude of the square of the mode’s coefficient. By definition the sum over the relative amplitudes of all modes is normalized to one at any given time, 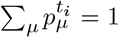. We order the modes by their relative amplitude from largest to smallest and take the cumulative sum of their values until a chosen cutoff percentage is reached, see Fig. 2A. We find that the percentage of modes required to reach a specified cutoff value varies considerably over time. For a 90 % amplitude cutoff we find that on average 6.06 % (≈ 70 modes) of the 1500 modes are significant with a large standard deviation of 36.96 %.

**Fig. 2.**
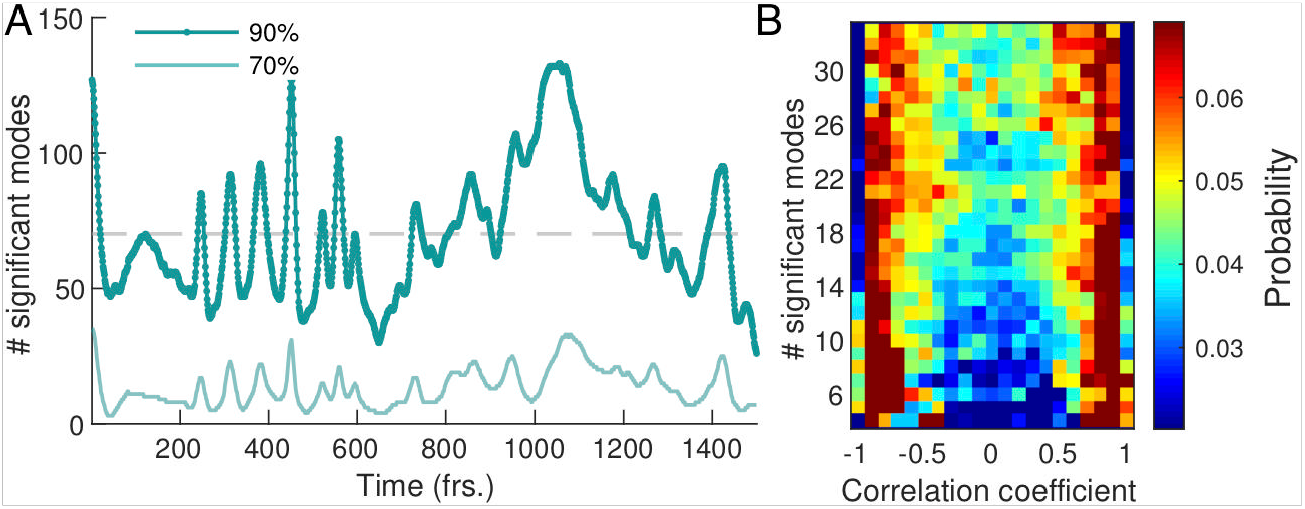
Dynamics of network contraction pattern is subject to strong variability in the percentage of significant modes and correlations between them. (A) Significant modes given by the percentage of the total number of modes required for the cumulative sum of their relative amplitudes to reach 70 % (light green) and 90 % (dark green) of the total amplitude plotted overtime. Gray dashed line indicates the 4.68% (≈ 70 modes) average of significant modes. (B) Distribution of coefficient correlation values depending on the number of significant modes taken from the 70 %-cutoff curve in (A). Correlation values show a trend from strong (anti-)correlation for a small number of significant modes (lower) to a more uniform distribution of correlation values for a large number of significant modes (upper).

Apart from the number of significant modes, the dynamics of the network depends on the temporal correlation of modes. In Fig. 2B we show the distribution of temporal correlations between mode coefficients as a function of the number of significant modes (see *Materials and Methods*). For a small number of significant modes the coefficients are strongly (anti-)correlated in time, while for a large number of significant modes, temporal correlations between coefficients are more uniformly distributed over the entire interval. The first is reflected in coordinated pumping behaviour/contractions, and the latter gives rise to irregular network-wide contractions. The above analysis shows that the dynamics of network contractions covers a wide range in complexity, from superposition of few large-scale modes strongly correlated in time, to superpositions of many modes of varying spatial scale and temporal correlations. This gives rise to strong variability in the regularity of the contraction dynamics over time. Up to now we investigated an ‘idle’ network not performing a specific task, so we next stimulate the network to provoke a specific behavior and scrutinize how the continuous spectrum of modes contributes to it.

### Stimulus response behavior is paired with activation of regular, large-scale contraction patterns interspersed by many-mode states

To probe the connection between a specific behavior and network contraction dynamics, we next apply a food stimulus to the same network, see Fig. 3A. Food acts as an attractant and causes locomotion of the organism toward the stimulus in the long term. The stimulus immediately triggers the network to inflate in a concentric region around the stimulus site. Also, the thick transport tubes oriented toward stimulus location increase their volume, see Fig. 3A. Altogether these morphological changes are typical for the specific behavior induced here, namely the generation of a new locomotion front.

**Fig. 3.**
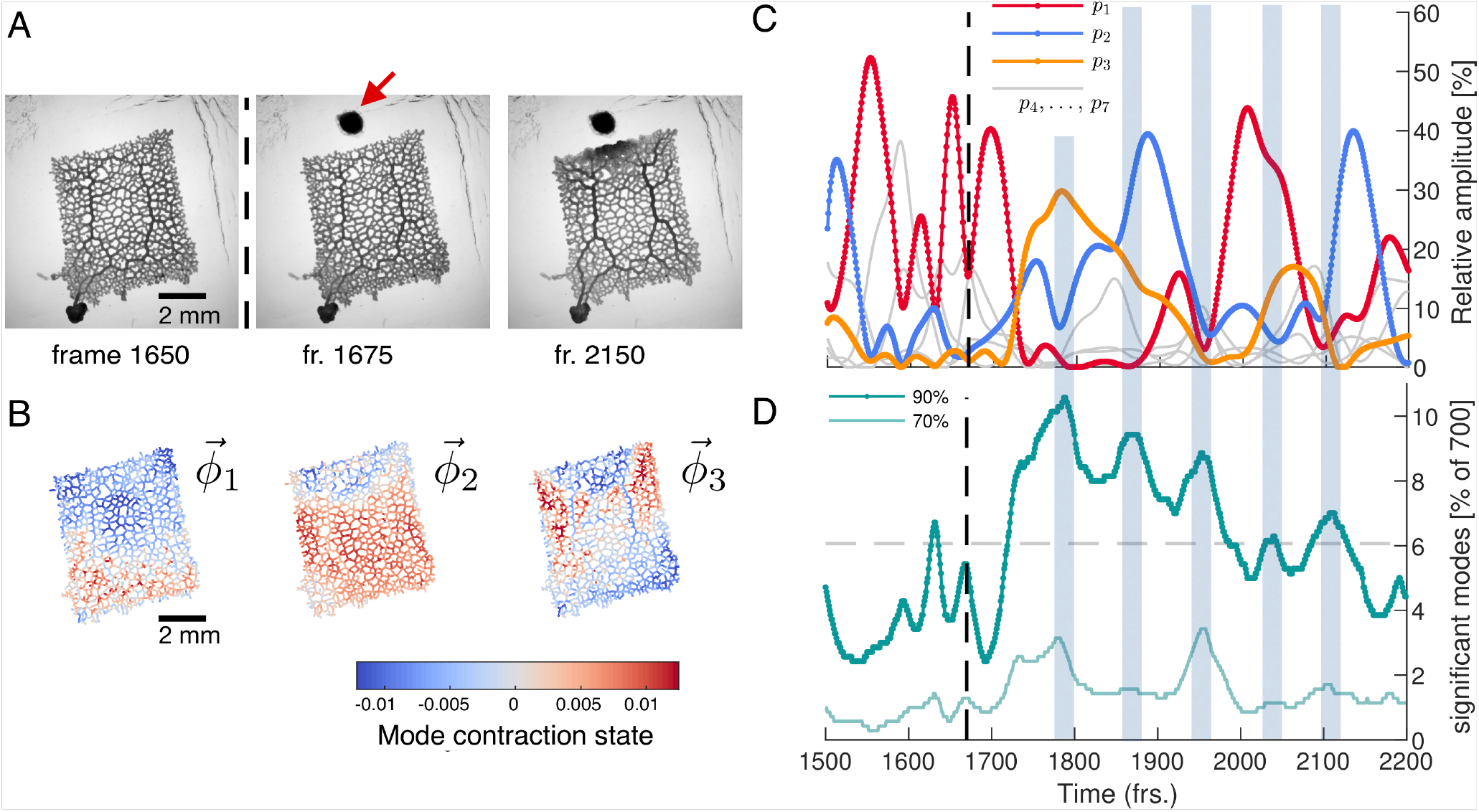
Network reacts to food stimulus with variations in the dynamics of the contraction pattern. (A) Bright-field frames showing the network’s growth response to a food stimulus (red arrow). (B) The spatial structure of the three top-ranked modes. (C) Dynamics of the relative amplitudes of the three top-ranked modes. After the stimulus (dashed line at frame 1670), time intervals with a single contraction mode dominating in amplitude (red, blue, yellow) over all other modes prevail. Mode amplitudes four to seven are shown in gray for reference. Interspersed we find time intervals where a larger number of modes have a similar amplitude. These times are indicated by the blue shaded boxes extending across (C) and (D). (D) Percentage of total number of modes required to reach a cumulatively summed amplitude of 70% (light green) and 70% (dark green) of the total amplitude, over time. Gray dashed line indicates the 6.06% (≈ 42 modes) average of significant modes.

To identify potential changes in the contraction dynamics due to stimulus application we perform PCA on a 700 frames long subset of the data subsequent to the data of the previous section. At first, we rediscover a continuous spectrum of modes displaying large variability in the number of significant modes resembling the ‘idle’ dynamic states, see *SI Appendix* Fig. S2. Yet, when mapping out the spatial structure of the highest-ranked contraction modes, see Fig. 3B, we discover that the now dominant modes clearly reflect the visually determined response of inflation close to the stimulus and the activation of the thick transport tubes. In fact, for more than 500 frames after the stimulus has been applied, the rhythmic contraction dynamics of the network are dominated by the three highest ranked modes, see Fig. 3C and *SI Appendix* Fig. S3 for the mode coefficients. During this period, every time a single mode is the most active one for a duration of > 30 frames, its amplitude exceeds that of any other mode by 20-30%. Taken together, this show that behavioral response, namely the generation of a new locomotion front, is linked to activation of distinct large-scale, regular contraction patterns.

Strikingly, these regular contractions dynamics are interspersed with many-mode states where the number of significant modes increases considerably, see Fig. 3D. The number of significant modes seems to oscillate after the stimulus and distinct maxima, of many modes, coincide with times at which the organism switches from one dominant contraction pattern to another, as indicated by the shaded regions in Fig. 3C and Fig. 3D. While our observations suggest that prolonged regular dynamics dominated by a few or even a single mode are associated with specific behavior like locomotion, the many-mode states seem to serve as transition states between them. While the network morphology is characteristic for *P. polycephalum*, reducing network complexity may help to conclude on the role of regular dynamics, many-mode states and the therefrom arising continuous distribution of modes.

### Number of significant modes determines maximum cytoplasmic flow rate in the minimal morphological representation of the network

We next perform exactly the same course of experiments as before but on a *P. polycephalum* specimen reduced in complexity to a single tube with a locomotion front at either end, see inset in Fig. 4. Strikingly, again we find a continuous spectrum of modes displaying a large variability in the number of significant modes, see *SI Appendix* Fig. S4 and S5. This observation finally underlines that the continuous spectrum of modes and its variability in activation is intrinsic to the organism’s behavior, ruling out that the complexity of contraction modes only mirrors morphological complexity. Foremost, this minimal constituent of a network allows us now to directly map the effect of variations in the contraction dynamics onto behavior. From the experimentally quantified tube contractions we calculate the maximal flow rate at any point of the tube (40) and over time correlate the strength of the flow rates, driving locomotion behavior at the tube ends, with the number of significant modes, see *SI Text*. For both the flow rate at the left and right end of the tube, shown in Fig. 4, and *SI Appendix* Fig S6, respectively, we find that large flow rates are only achieved when the number of significant modes is small. We had previously found that few significant modes are highly (anti-)correlated, whereas states with many significant modes are not, see Fig. 2B. This observation now confirms our physical intuition that the irregularity of states consisting of many modes goes hand in hand with reduced pumping efficiency and thus unspecific behavior. Since a small number of significant modes not necessarily always implies a large flow rate, we next turn to analyze their exact spatial structure and in-stantaneous temporal correlation to determine how cytoplasmic flow rates impact behavior.

**Fig. 4.**
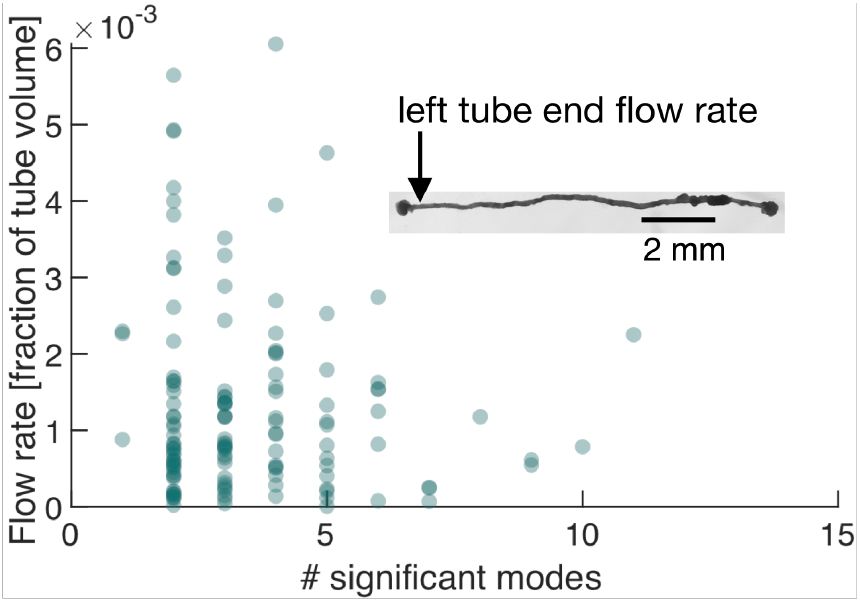
Number of significant modes is indicative for the volume flow rate in a cell reduced in its network complexity to a single tube. Inset: Single tube with locomotion fronts at both ends. Main plot: Volume flow rate at the left tube end, calculated from tube contraction dynamics versus the number of significant modes at different times. High flow rates are only achieved for a small number of significant modes.

### Instantaneous coupling and selective activation of modes determine locomotion behavior

We now demonstrate the impact of changes in the dynamics of a small number of modes on the organism’s behavior. For this we quantify the locomotion behavior of the single tube by tracking the area of the locomotion fronts protruding from each end of the tube over time, see Fig. 5B(ii). While initially the right end is protruding faster at the expense of the left end, a food stimulus applied to the left end of the tube flips the direction of locomotion.

**Fig. 5.**
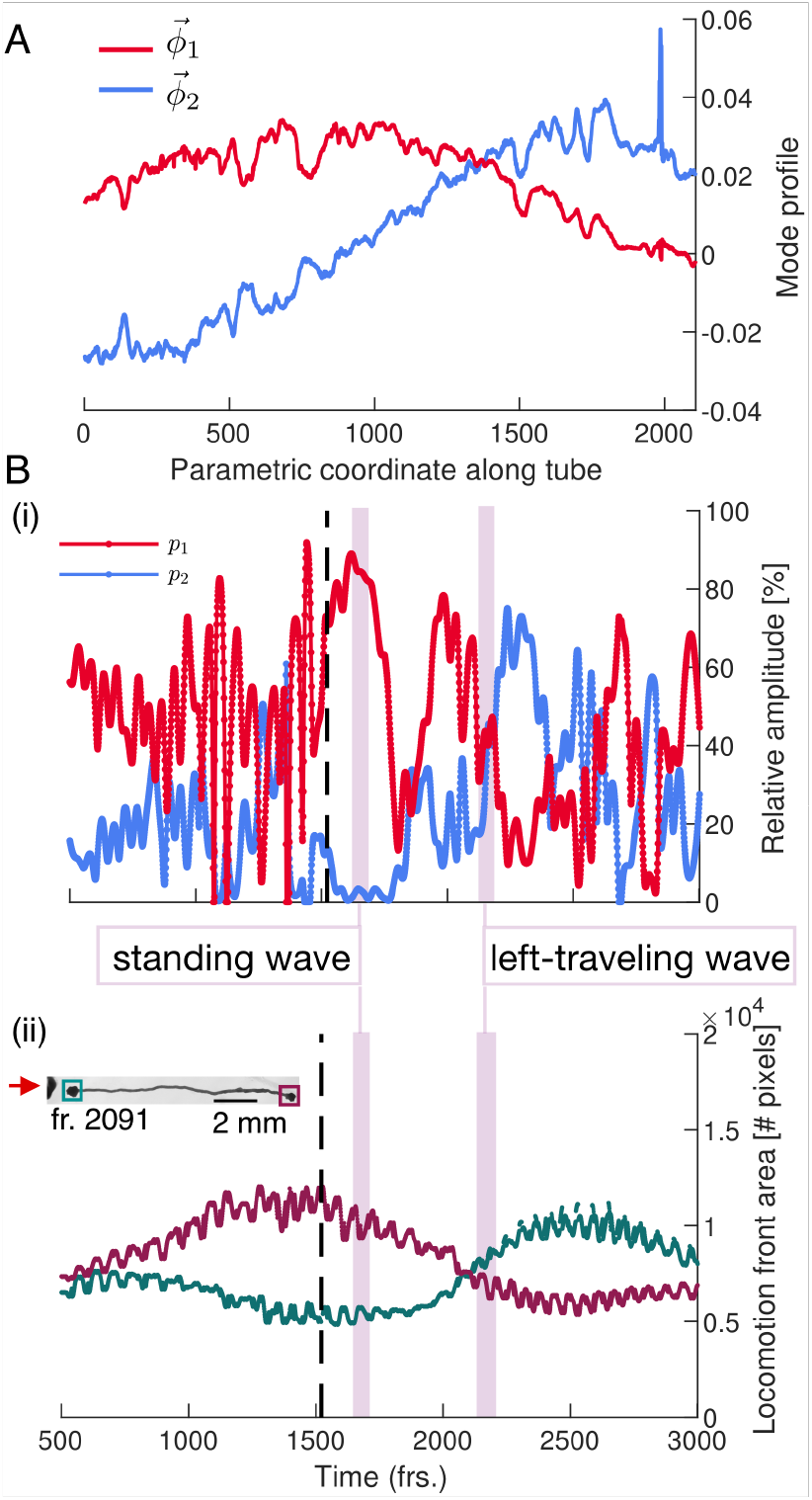
Locomotion behavior of a single tube is determined by activation and temporal coupling of modes. (A) The two top-ranked modes. (B) (i) Activation of the two top-ranked modes given by their relative amplitude. (ii) Behavior of the locomotion front at each end of the tube over time. Vertical pink bars indicate two representative time intervals and the nature of the two-mode superposition is specified.

Turning to the modes that drive this behavior, we find that over long time intervals, and in particular after the stimulus, the two top-ranked modes dominate the tube’s contraction dynamics, see Fig. 5(B)(i) and *SI Appendix* Fig. S5 and S7. By parametrizing the single tube with a longitudinal coordinate, the spatial shapes of these dominating modes 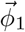 and 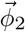 approximate Fourier modes, see Fig. 5(A). To illustrate the connection between tube contraction dynamics and locomotion behavior, we pick two representative time intervals after the stimulus where either only mode 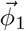 or modes 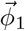 and 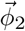 equally dominate overall, see vertical pink bars in Fig. 5(B). During the first interval when mode 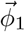 alone is dominating, the tube is driven by a standing wave contraction pattern – yielding only a low cytoplasmic flow rate. Correspondingly, the size of the locomotion front at either end shows no significant change in area during this interval. In contrast, during the interval when both modes 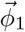 and 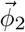 are equally active, the resulting superposition is a left-traveling wave producing a large cytoplasmic flow rate in that direction. The left-traveling wave is in accordance with the growth of the left and retraction of the right locomotion front. These two examples solve the conundrum of Fig. 4, which shows that a small number of significant modes does not necessarily lead to high cytoplasmic flow rates. Yet, the direct mapping of contraction dynamics onto ensuing cytoplasmic flows confirms that a small number of significant modes is associated with specific behavior. High cytoplasmic flow rates at the tube ends drive locomotion, while lower flow rates likely lead to other behaviors such as mixing. Furthermore, many-mode states seem necessary for transitions in a multi-behavioral space.

## Discussion

To uncover the origin of behavior in *P. polycephalum* we quantified the dynamics of living matter and linked it to emerging behavior. The simple build of this non-neural organism allows us to trace contractions of the actomyosin-lined tubes, compute cytoplasmic flows from the contractions and finally link these dynamics to the emerging mass redistribution and whole-organism locomotion behavior. Decomposing the contractions across the network into individual modes, we discover a large intrinsic variability in the number of significant modes over time along a continuous spectrum of modes. Triggering locomotion, we identify that states with few significant modes and regular contraction patterns correspond to specific behaviors like locomotion. Yet, also irregular contraction patterns consisting of a large number of significant modes are present, particularly marking the transitions between different regular states. Our findings suggest that a continuous spectrum of contraction modes allows the living matter network *P. polycephalum* to quickly transition between a multitude of behaviors using the superposition of multiple contraction patterns.

Our observation of interlaced regular and irregular contraction patterns in *P. polycephalum* reminds of the strongly correlated or random firing patterns of neurons in higher organisms (3). Also in neural organisms, stereotyped behaviors are associated with controlled neural activity, as for example for locomotion in *C. elegans* (41) or the behavioral states of the fruit fly *Drosophila melanogaster* (42, 43). Though rarely discussed, variability in the dynamics of behavior is also observed in such neuronal organisms (44, 45). It is, thus, likely that the transition role of irregular states consisting of many significant modes observed here for *P. polycephalum* parallels the mechanisms of generating behavior in higher forms of life.

*P. polycephalum* is renowned for its ability to make informed decisions and navigate a complex environment (11–17, 46, 47). It would be fascinating to next follow the variability of contraction dynamics during more complex decision-making processes. Further, it would be interesting to observe ‘idle’ networks during foraging over tens of hours. It is likely that the contraction states with many significant modes here act as noise that can spontaneously cause the organism to ‘decide’ and reorient its direction of locomotion.

*P. polycephalum’*s body-plan as a fluid-filled living network with emerging behavior finds its theoretical counterpart in theories for active flow networks developed recently (48, 49). Strikingly, these theories predict selective activation of thick tubes which we observe in the living network as well, prominently appearing among the top ranking modes, see 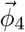 Fig. 1C(i) or 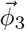 in Fig. 3C. This is a first hint that dynamics states arising from first principles in active flow networks could map onto behavioral and transition states observed here.

Likely our most broadly relevant finding in this work is that irregular dynamics, here arising in states with many significant modes, play an important role in switching between behaviors. This should inspire theoretical investigations to embrace irregularities rather than focusing solely on regular dynamic states. The most powerful aspect of *P. polycephalum* as a model organism of behavior lies in the direct link between actomyosin contractions, resulting in cytoplasmic flows and emerging behaviors. The broad understanding of the theory of active contractions (30–34) might therefore well be the foundation to formulate the physics of behavior not only in *P. polycephalum* but also in other simple organism. This would not only open up an new perspective on life but also guide the design of bio-inspired soft robots with a behavioral repertoire comparable to higher organisms.

## Methods

### Experiments

The specimen was prepared from fused microplasmodia grown in a liquid culture (50) and plated on 1,5%-agar. The network was trimmed and imaged in the bright field setting in Zeiss ZEN 2 imaging software with a Zeiss Axio Zoom V.16 microscope equipped with a Hamamatsu ORCA-Flash 4.0 digital camera and Zeiss PlanNeoFluar 1x/0.25 objective. The acquisition frame rate was 3 sec. The stimulus was applied in a form of a heat-killed HB101 bacterial pellet in close network proximity.

### Distribution of temporal correlations

For a given time point *t_i_*, the significant modes are determined based on the 70% criterion curve from Fig. 2A. Next, the temporal correlations among the coefficients are computed in a time interval of ±15 frames around the time point *t_i_*. The correlations are then counted in bins of the appropriate row of Fig. 2A. Repeating this processing for all time points and normalizing each row by the total number of correlations in that row, we obtain the final distribution shown.

## Acknowledgements

This work was supported by the Max Planck Society. In the final stages of this work P.F. was supported by the Simons Foundation program for Mathematical Modeling of Living Systems, grant number 400425. M.K. acknowledges the support of IMPRS for Physics of Biological and Complex Systems.

