## Supplemental Material for "Emergence of behavior in a self-organized living matter network"

### Data

The typical thickness of tubes in a *P. polycephalum* network is $\sim$ 50-100 µm and the contraction amplitude about $\sim$10 % of the tube's typical thickness [1]. This change in tube thickness can be detected from a bright-field microscopy recording. We record one bright-field frame every three seconds. Since the periodic contractions of the tubes take place on the time scale of 100 sec, they are thus well resolved by the selected frame rate. Typically an idle network keeps a stable morphology and does not move significantly over a period of 1.5 h to 2.5 h which we use for recording its contraction dynamics.

### Data processing

Our data is a stack of bright-field images recorded from the *P. polycephalum* network with a rate of one frame every three seconds. Each bright-field frame has a time label $t_i$ and the total number of frames is given by $T$. We process this data in the following steps. First, we mask the network in the bright-field images through thresholding. It is important to note that we use the same mask for all the images in the stack. This is possible since we consider a network that does not significantly move or change its morphology. This is true even when we apply a stimulus to the network, since we only consider the initial stages of stimulus response, before the network starts to display strong movement. From the masked regions of the bright-field frames, we extract pixel intensity values which we convert to 8-bit format. Since we are here primarily interested in the contraction dynamics of the organism and not in the the actual base thickness of tubes or its long-term growth dynamics, we detrend the data using a moving-average filter (rational transfer function) with a window size of two contraction periods (∼200 sec) [2]. This leaves us only with the desired information about contractions taking place in the time scale of several minutes. We store the intensity values of each frame in a vector $\vec{I}^{t_i}$ of dimension $M$ equal to the number of pixels in the network, and $i$ indexes the frames in the range $i = 1, \ldots, T$. From the post-processed data we define the following data matrix

$$\mathbf{X}^t = \left( \vec{I}^{t_1}, \vec{I}^{t_2}, \ldots, \vec{I}^{t_T} \right) , \tag{1}$$

where $t$ denotes the matrix transpose.

### Principal Component Analysis (PCA)

The contraction modes are computed from the covariance matrix of the data. We compute the covariance matrix from the data matrix $\mathbf{X}$ after subtracting the mean from each column. The covariance matrix is given by

$$\mathbf{C} = \frac{1}{T-1} \mathbf{X}^t \mathbf{X} \, . \tag{2}$$

The sought after contraction modes $\vec{\phi}_\mu$ are the eigenvectors of the covariance matrix

$$\mathbf{C} \vec{\phi}_\mu = \lambda_\mu \vec{\phi}_\mu \, , \tag{3}$$

and $\lambda_\mu$ is the eigenvalue. The number of non-zero eigenvalues is equal to the rank of the covariance matrix. The eigenvalue captures the variance of the data along the direction of mode $\vec{\phi}_\mu$. We also define the relative eigenvalue as

$$\tilde{\lambda}_\mu = \frac{\lambda_\mu}{\sum_{\nu=1}^{T} \lambda_\nu} \, . \tag{4}$$

The mode coefficient $a_\mu$ is obtained by projecting the data onto mode $\vec{\phi}_\mu$.

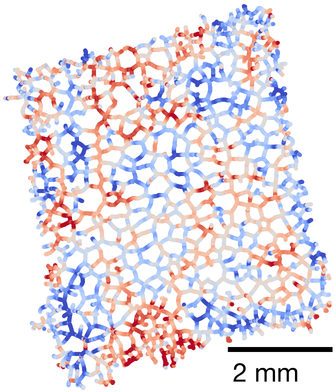


Figure S1: Contraction mode $\vec{\phi}_{30}$ obtained from Principal Component Analysis of the large rectangular network.

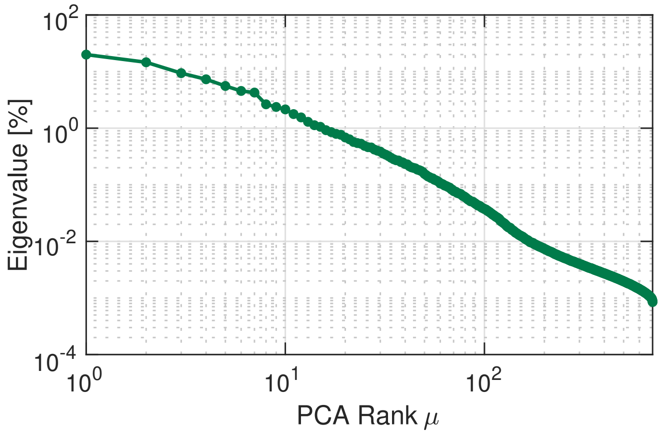


Figure S2: Principal Component Analysis yields a continuous spectrum of contraction modes in the *P. polycephalum* network exposed to a food stimulus. The eigenvalue spectrum shown here is computed from data consisting of 700 frames.

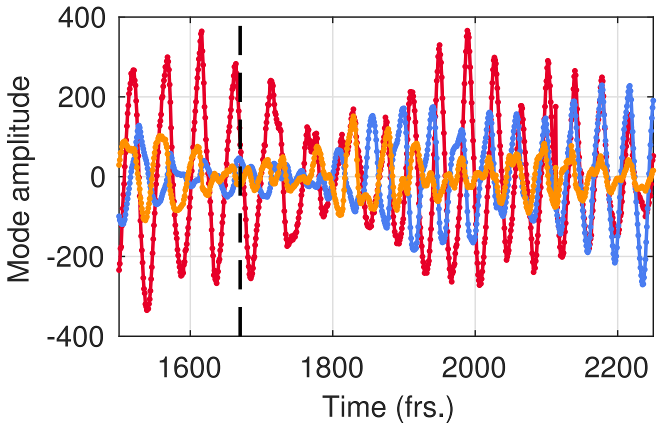


Figure S3: Temporal dynamics of the coefficients of the three top-ranked contraction modes $\vec{\phi}_1$ (red), $\vec{\phi}_2$ (blue) and $\vec{\phi}_3$ (yellow) in the large rectangular network. The variation in amplitude, frequency and relative phase shift between coefficients is captured before and after application of the stimulus to the network. The time of stimulus application is indicated by the dashed line.

### Flow rate calculation in a *P. polycephalum* cell with single-tube morphology

To compute the flow rate of the cytoplasm in a *P. polycephalum* specimen with single-tube morphology we use the theory developed in [3, 4]. In that work the flow of an incompressible Newtonian fluid inside an axisymmetric tube of fixed length is considered and the equations for the flow velocity field are written in the lubrication theory approximation. Furthermore, a time-dependent thickness profile of longitudinal waves is imposed in the tube. Assuming no-slip boundary conditions, the flow field can be fully determined at every point along the tube as a function of the time-dependent tube profile. For the case when the tube profile is a periodic train of waves, we compute the volume flow rate averaged over an oscillation period by evaluating equation 13 of [4]. We express the flow rate in units of volume of the entire tube divided by the oscillation period. This serves to characterize the performance in pumping of the significantly contracting *P. polycephalum* cell. We determine the time period over which to average the volume flow rate directly from the flow rate curve. Furthermore, the thickness profile of the tube is given by the measured pixel intensity profile.

### Mode superpositions in a *P. polycephalum* cell with single-tube morphology

We are interested in how the contraction dynamics of the cell controls the cell's locomotion behaviour. In our analysis we therefore focus on the tube segment connecting the locomotion fronts at either end of the tube and perform Principal Component Analysis only on this part of the cell. Since the tube is effectively one-dimensional, we find that the modes we obtain closely approximate Fourier modes. This means that superpositions of these modes afford a clear interpretation in terms of different contraction wave patterns. Such an interpretation is even further facilitated by the fact that we find that over large time intervals after the stimulus, the number of significant modes is very small. Indeed, over such time intervals it is sufficient to approximate the contraction dynamics with only one or two modes, as can be seen from Figs. S5 and S7. Hence we are essentially studying a superposition of modes $\vec{\phi}_1$ and $\vec{\phi}_2$ shown in Fig. 5A of the main text with their oscillating mode coefficients shown in Fig. S8. To develop a more intuitive understanding of the nature of the superposition, we note that the modes $\vec{\phi}_1$ and $\vec{\phi}_2$ approximate sine and cosine functions over the length of the tube. Given a sine and cosine spatial contraction profile different types of superpositions can be formed depending on the nature of their time-dependent coefficients. To illustrate further let us assume the idealised case where both coefficients are sine functions that can have differing phases and amplitudes. Then, if the coefficient of one contraction profile is very small compared to the other, the resulting superposition is a standing wave. In case the coefficients have equal amplitudes, but are phase shifted by $\pi/2$, the superposition is a traveling wave. Finally, if the coefficient amplitudes are not equal and the phase shift lies somewhere between zero and $\pi/2$, the nature of the superposition is a mix of standing and traveling wave. Extrapolating this idealised picture allows us to infer the contraction dynamics resulting from our two-mode approximation. We see that the coefficients of the two modes $\vec{\phi}_1$ and $\vec{\phi}_2$ shown in Fig. S8 change in amplitude and phase relative to each other. It is easy to identify from this plot together with the plot if relative amplitudes in Fig. S7 time intervals which approximate one of contraction dynamics that we have described for the idealised system. Therefore we conclude that the superposition of the two top modes changes in its nature over time, ranging form a purely standing wave to a pure traveling wave.

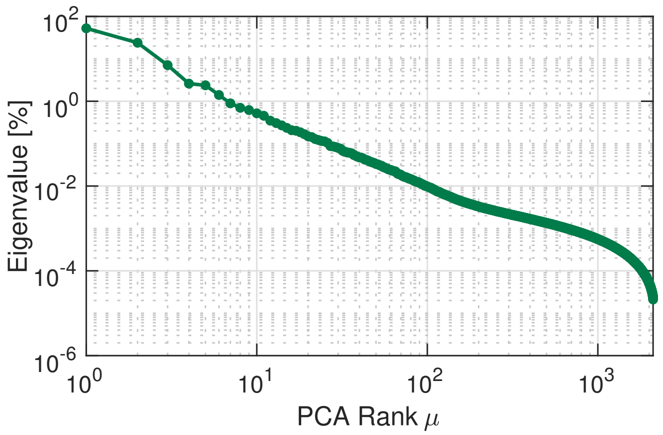


Figure S4: Principal Component Analysis yields a continuous spectrum of contraction modes in a *P. polycephalum* specimen with single-tube morphology. The eigenvalue spectrum is computed from a data segment which is 3000 frames long and includes the time when a food stimulus is applied to one end of the tube.

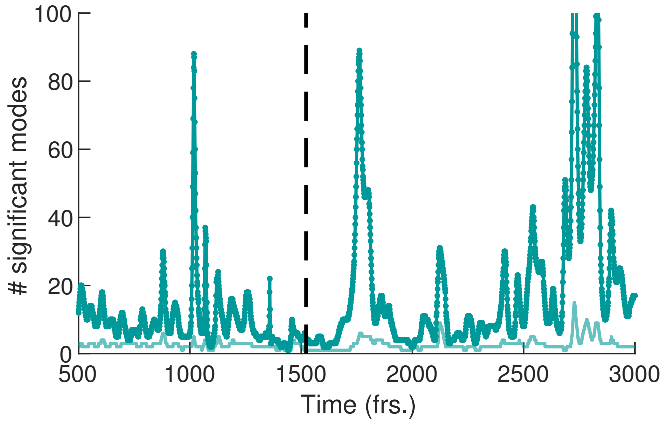


Figure S5: Temporal variation in the number of significant modes for the single-tube *P. polycephalum* network. Number of significant modes is shown for a cutoff criterion of 70% (light green) and 90% (dark green) of the total relative contraction amplitude. The time of stimulus application is indicated by the dashed line.

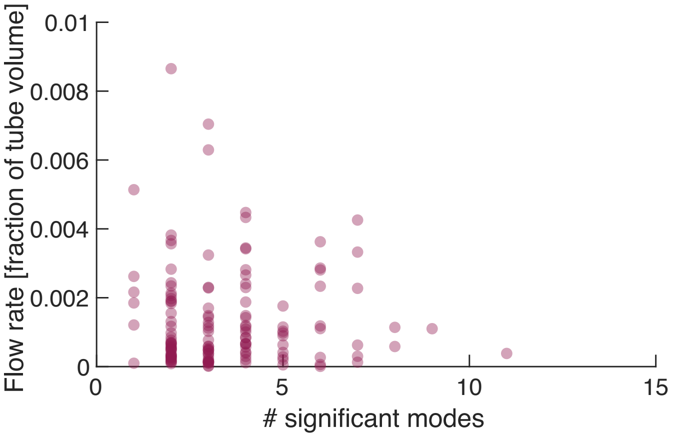


Figure S6: Number of significant modes is indicative for the volume flow rate in a cell reduced in its network complexity to a single tube. Main plot: Volume flow rate at the left tube end, calculated from tube contraction dynamics versus the number of significant modes at different times. High flow rates are only achieved for a small number of significant modes.

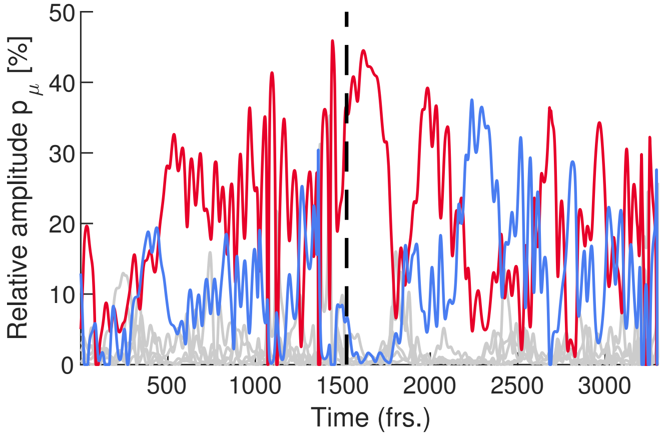


Figure S7: Comparison of the activity of top-ranked modes of the single-tube *P. polycephalum* network as quantified by their relative amplitude. We show the relative amplitude of mode of mode $\vec{\phi}_1$ (red), mode $\vec{\phi}_2$ (blue) and the significantly smaller relative amplitudes of modes $\vec{\phi}_3$ to $\vec{\phi}_7$ are shown in gray.

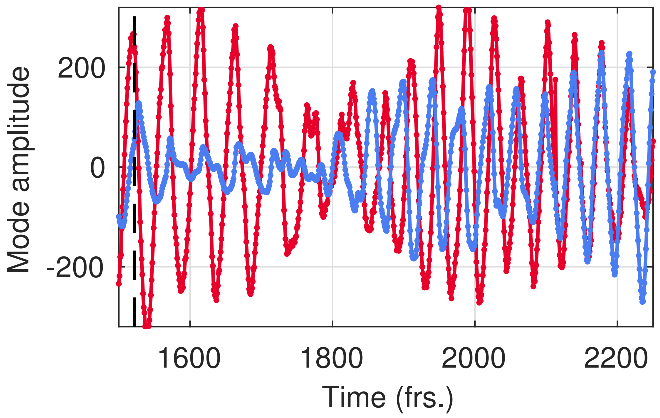


Figure S8: Temporal dynamics of the coefficients of the two top-ranked contraction modes $\vec{\phi}_1$ (red) and $\vec{\phi}_2$ (blue) in the single-tube *P. polycephalum* network. The variation in amplitude, frequency and relative phase shift between coefficients is captured shortly before and after application of the stimulus to the single tube. The time of stimulus application is indicated by the dashed line.
